# Nestling-Care Decisions by Cooperatively Breeding American Crows

**DOI:** 10.1101/2024.02.01.578448

**Authors:** Carolee Caffrey, Charles C. Peterson, Tiffany W. Hackler

**Author notes:** 5235 Kester Ave #318 Sherman Oaks CA 91411. Copper Mountain College, Joshua Tree CA 92252. 8924 S. Old Dutch Rd, Rogers, AR 72756, USA.

## Abstract

During the nestling stage of breeding seasons in Stillwater, OK, pairs of American Crows (*Corvus brachyrhynchos brachyrhynchos*) lived alone or in groups of variable composition; auxiliaries included individuals that had delayed dispersal, immigrated into groups, or returned to natal territories after having lived elsewhere. Most, but not all, auxiliaries contributed to feeding nestlings, and their contributions varied considerably. On average, breeders fed nestlings at greater rates than did auxiliaries, and female breeders spent more time at nests than did other group members. Breeders compensated for auxiliary contributions by reducing their own; this and breeder responses to the disappearance of auxiliary feeding group members provide evidence that these long-lived, iteroparous animals were managing energy budgets so as to maximize fitness over the long term. Female breeders in larger groups spent more time at nests than did those in smaller groups, but not for expected reasons and not to any reproductive benefit. A few female auxiliaries spent increasing amounts of time at nests as nestlings aged. No other measured phenotypic characteristic of individuals was found to explain any of the wide variation in the patterns of nestling care exhibited by members of our population.

## INTRODUCTION

In the literature of cooperative breeding exist many documentations of the contributions of group members to nestling feeding, with statistical analyses of variables possibly influencing individual decisions. Theoretical treatments of empirical data have considered decisions made by group members in such contexts as whether to breed or help (e.g., Emlen and Wrege 1991, Wrege and Emlen 1994), whether to help or not (e.g., Marzluff and Balda 1990, Dickinson and Hatchwell 2004, Hatchwell et al. 2013), whom to help (e.g., Emlen and Wrege 1988, Russell and Hatchwell 2001), whom to harass (e.g., Emlen and Wrege 1992), whether to compensate in response to helper contributions or to use those contributions additionally (e.g., Hatchwell 1999, Heinsohn 2004, Carranza et al. 2008), and how to respond to the disappearance of other group members (e.g., Baglione et al. 2010, Liebl et al. 2016a and b).

In the above and other such studies of proximate cues and/or ultimate explanations for helping behavior, it has been customary to ignore individual variation and use means to describe group members of a particular status - breeding females, adult male auxiliaries, or group members related to breeders by particular values, e.g. - as if all individuals of a particular status were the same. This pattern has been typical of the field of behavioral ecology overall; the tendency to seek adaptive optima and view individual variation as nonadaptive “noise” has been acknowledged by authors for decades (e.g., Hirsch 1963, Clark and Ehlinger 1987, Wilson 1998, Sih et al. 2004, Dall et al. 2004, Réale et al. 2007, Carter et al. 2013). Acknowledged also for decades (e.g., Slater 1981, Clark and Ehlinger 1987, Wilson 1998, Bolnick et al. 2003) and gaining increased support, particularly recently and in conjunction with “personality” thinking, is the likelihood that the variable strategies being pursued by individual group members may themselves be adaptive (e.g., Réale et al. 2010, Careau and Garland 2012, Wolf and Weissing 2012 and references therein, Dingemanse and Araya-Ajoy 2015 and references therein).

American Crows in Stillwater, OK, bred cooperatively (Caffrey and Peterson 2015). Individuals in this population made variable residency decisions, and many unpaired individuals moved in and out of groups throughout the year (Caffrey and Peterson 2015). A common time to disperse out of groups was during the week or so preceding hatching: 29% of auxiliaries present during nest building and incubation (“pre-hatch auxiliaries”) moved out of groups for nestling and post-fledging periods (most returned later in the year; Caffrey and Peterson 2015). For breeders and the diverse auxiliaries who chose to remain for nestling-rearing periods, we investigated a question rarely addressed by others (Cant and Field 2001, Heinsohn 2004, Eguchi et al. 2009, Bergmüller et al. 2010, Koenig and Walters 2012; surprising as helping behavior in most species is facultative): Once individuals have chosen to contribute to nestling care, how do they next decide how much to invest in feeding? In view of our results, we hereby add our voices to the chorus calling for increased focus on the existence, causes, and consequences of variation in behavior among individuals (e.g., Komdeur 2006, Bergmüller et al. 2010, Carere and Locurto 2010, Careau and Garland 2012, Dingemanse and Araya-Ajoy 2014, Webster and Ward 2011).

## METHODS

### Background

Our study population in Stillwater, Oklahoma, had been under observation since August 1997 (Caffrey and Peterson 2015). Individual crows were captured as free-flyers (Caffrey 2002a) or obtained as nestlings, and marked with patagial tags and unique combinations of colored plastic and U.S. Fish and Wildlife Service metal leg bands (Caffrey 2002b and c). Crows caught as free-flyers were aged via inspection of plumage and mouth color pattern characteristics (Emlen 1936); adults were assigned the age “≥2 y” at marking. For the present study, crows were observed during the nestling stages of the breeding seasons of 2001 and 2002; late March through early June in both years. Observations were made with binoculars and spotting scopes, primarily from vehicles, except for one group in 2001 (#29: Table 1 and Appendix 3b [and Table 1 in Caffrey et al. 2016a, and Tables 1 and 3, and Appendix A 1a, in Caffrey and Peterson 2015]); this wary groups’ placement of their nest next to a window in an empty building allowed continuous videotaping of nest activities for up to 10 hours at a time on many post-hatch days (on others - days 19-24 and 31-35 – we were locked out of the building and unable to observe Group 29).

**Table 1.**
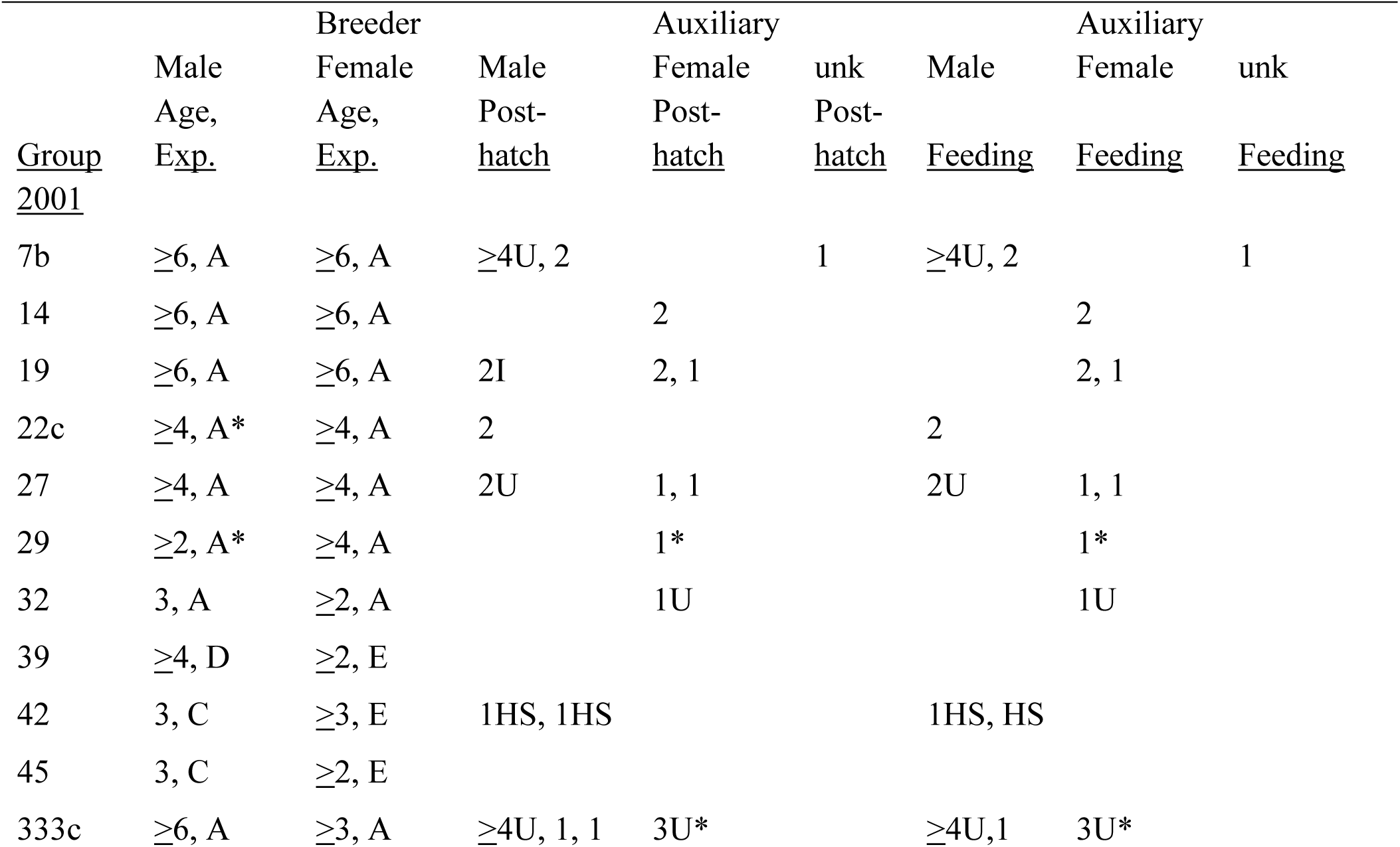
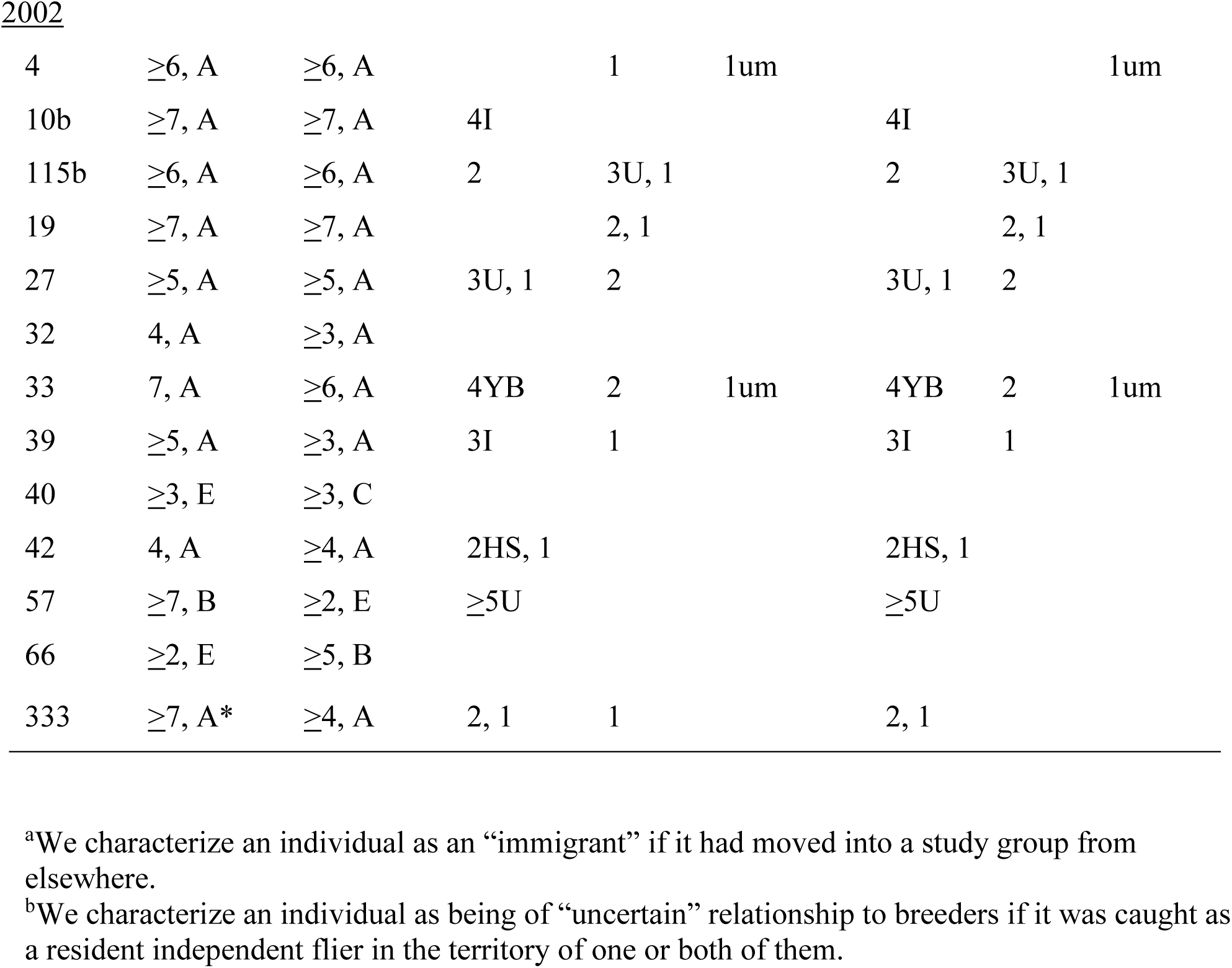
Composition of crow Post-hatch and Feeding Groups. Each numeral represents an individual by its age in years. Experience (Exp.): A = experienced/nested with current mate in the previous year, B = experienced/did not nest with current mate in previous year (#66 in 2002) or attempt (#57 in 2002; Appendix A 2c in Caffrey and Peterson 2015, and Appendix 2a and Fig. 1g in Caffrey et al. 2016a), C = nesting for first time, D = likely nesting for first time, E = history unknown. Most auxiliaries had hatched in nests of one or both breeders. I: immigrant^a^. HS: social half-sib of male breeder. YB: younger brother of male breeder. U: dispersal history (relative to group) uncertain^b^. um: unmarked. unk: sex unknown. Asterisks indicate individuals that disappeared from post-hatch and feeding groups.

### Group Composition

We examined the nestling-care decisions made by auxiliaries who chose to remain with pre-hatch groups for nestling-rearing stages of breeder reproductive attempts. We defined auxiliary members of post-hatch groups as individuals observed with at least one breeder in at least 35% of all observations of breeders away from nest areas. Feeding groups included all post-hatch group members observed to make at least one feeding trip during nestling feeding watches (below). In Table 1, we distinguish between auxiliaries hatched in nests of one or both breeders, auxiliaries that immigrated into groups, and auxiliaries originally caught (and marked) as independent flyers in territories of groups (and so whose dispersal histories, relative to breeders, were uncertain). We later determined parentage for six of the seven individuals of uncertain relationship to breeders (eight auxiliaries in Table 1); see below. Hereafter we use such terms as “mother,” “daughter,” “brother,” and “offspring” when the genetic parentage of particular individuals is known, and “social daughter” and “social son” for auxiliaries marked in nests of breeders (social mothers and fathers) but for whom parentage analysis was not performed. The three half-sibs of the male breeder in Table 1 (Group 42) were individuals with the same father but a different mother.

### Genetic Sampling and Analyses

We collected approximately 1cc of blood from the brachial vein of each crow for molecular determination of sex and for estimation of microsatellite-based relatedness values, analyzed parentage in the program Cervus (Kalinowski et al. 2007), and estimated microsatellite-based relatedness coefficients (r) in the program Relatedness (Queller and Goodnight 1989); details in Caffrey and Peterson (2015).

We did not have DNA samples for four individuals: the adult female auxiliary of uncertain relationship to breeders in 2001 (Group 333, “3U”), and the three auxiliaries whose sexes were unknown (Table 1). Throughout this paper, where appropriate, we report r values as means ± 95% CIs.

### Individual Contributions to Nestling Care

We made detailed observations of the nest-associated activities of individual crows. We identify first, second, and third nesting attempts (within years) as “a,” “b,” and “c.”

Hatch dates were determined primarily via nest watches and were identified as the first days feeding trips were observed (changes in female breeder and auxiliary behavior also signaled hatching had begun: below, Results, Hatching and Nestling Periods). There were four cases where nest watches had not been done in the 2-3 days prior to the first observed feeding trip; for these groups we estimated hatch dates based on crow behavior during brief checks during those days and during nest watches in the days thereafter. Nests were monitored until attempts were abandoned or until the day the last nestling fledged. We counted as fledged any young of the year observed alive outside its nest.

For groups other than #29 (above; Background), we conducted at least three nestling-feeding watches per week. For Group 29, we videotaped nine watches from Day 0 post hatching through Day 18, and five watches from Day 25 through 30, for a total of 5224 minutes (upon getting back into the building on Day 36, three of four nestlings had already fledged). During feeding watches, we recorded the behavior performed and amount of time spent at nests per visit by each individual. A feeding trip was attributed to any individual arriving at a nest with food that was fed to nestlings directly or via another group member. We included in analyses data for only one attempt per group per year. We conducted 151 total feeding watches at 11 nests in 2001 (including Group 29; mean = 188.8 min), and 188 at 13 nests in 2002 (mean = 96.5 min).

We examined the potential effects of many variables on the feeding contributions of group members. For all individuals, included were sex, post-hatch group size, feeding group size, nesting attempt (a, b, or c), and average genetic relatedness to nestlings. We were unable to determine many clutch or early brood sizes (most nests were difficult to access and crows often abandon nesting attempts upon disturbance), but for individuals in groups wherein nestlings had been marked, we also included total brood mass at marking and mean nestling “condition” at marking (residuals from linear regression of log mass on log tarsus length) as possible covariates of feeding effort. For individuals in groups for which the number of fledglings could be accurately determined, we included that measure of nesting success as well.

For breeders, we additionally examined the potential effect of prior experience, comparing individuals a) breeding for the first time, b) experienced and breeding with the same mates as in preceding attempts, and c) experienced and breeding with individuals other than mates in preceding attempts. For auxiliaries, we additionally examined potential effects of age, relationship with breeders (nestling of one, or both, or genetic offspring of one, or both, or not: brother, half-sib, or immigrant [Table 1]), as well as r values with breeders of both sexes.

Data from nest watches were statistically analyzed for patterns related to individual- and nest-level variables with use of SYSTAT’s General Linear Model module. Rates of nestling feeding changed over the time course of nestling periods (Fig. 1a and b), and therefore a potential source of bias was the inevitably different timing of watches across nests. Because some individuals contributed to feeding for only part of the nestling period (e.g., in cases of the disappearance of individuals or nest failure during nestling periods), direct comparison of feeding rates would be confounded by such unequal temporal sampling. In an attempt to remove this potential source of bias, we constructed overall quadratic regressions of behavioral rates on day of nestling period and its square, and then calculated residual rates from the equation for each individual during each watch. Individuals (and, analogously, nests) were then represented by their average residuals over their entire sampling period for statistical analyses.

**Fig. 1.**
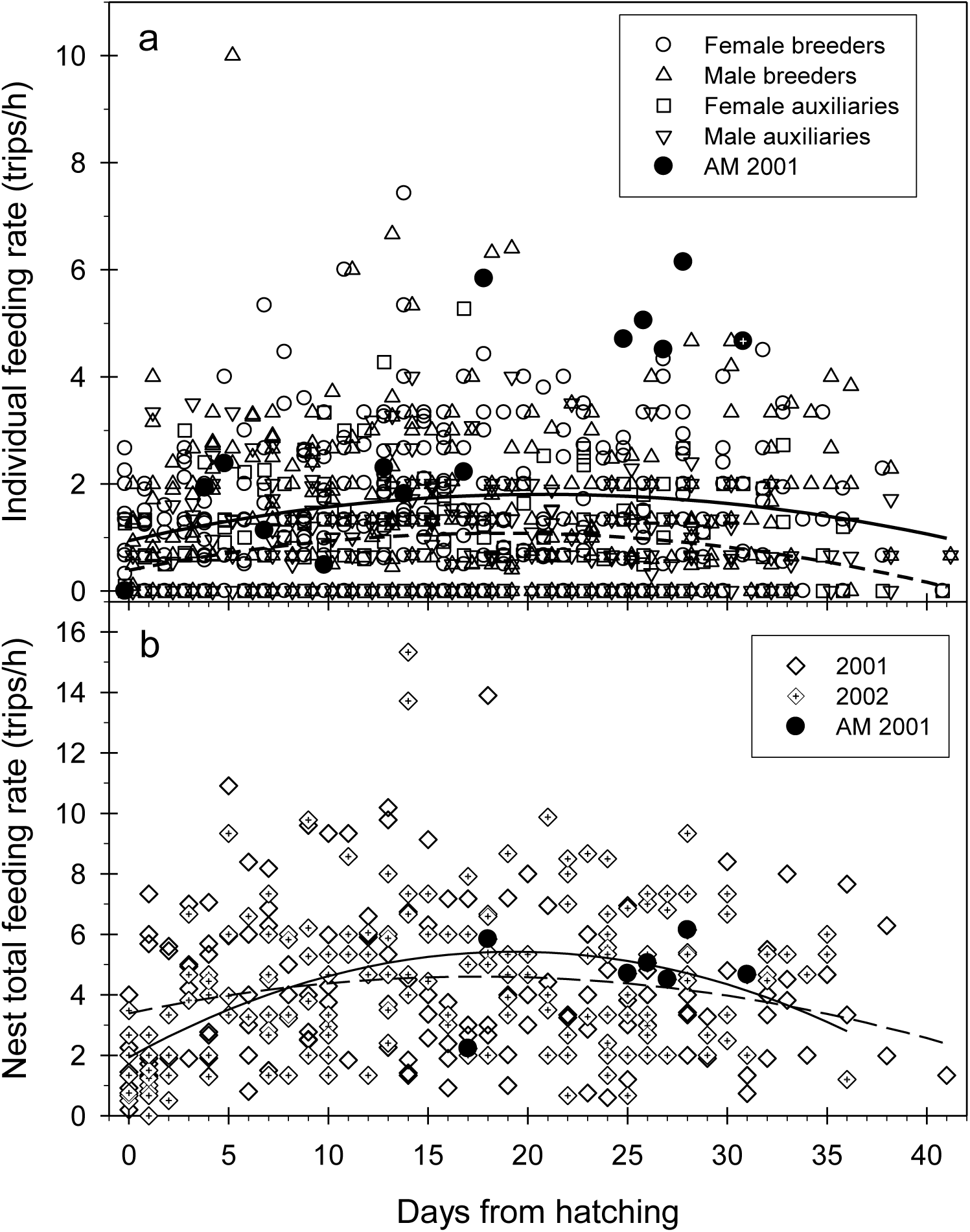
Nestling feeding rates in 2001 and 2002. (a) Trips/hr by individual group members. Solid curve is best-fit quadratic for breeders; dashed is that for auxiliaries. (b) Nest-wise trips/hr to nests. Solid curve is 2001, dashed is 2002. AM had been rendered the sole member of her group (see text and Appendix 3b). The crossed symbol for AM represents a watch after food was supplied.

We treated nesting attempts in different years as independent. A variety of preliminary analyses showed no differences between years and so data from the two years of this study were combined for statistical analyses of behavioral rates and time spent at nests.

Our field data were characterized by wide variation (Fig. 1), and our purposes in testing statistical hypotheses are largely heuristic. We therefore apply a liberal criterion of statistical significance, discussing all results with a probability of Type I error (P) ≤ 0.1.

## RESULTS

### Post-hatch and Feeding Group Composition

In 2001 and 2002, respectively, 77% of 11, and 82% of 13 post-hatch and feeding groups included at least one auxiliary. Post-hatch group size ranged from 2 to 6 with means of 3.7 in both years; feeding groups contained 2 to 5 individuals with average sizes of 3.6 and 3.5 in 2001 and 2002, respectively (Table 1). There was no sex bias to the post-hatch auxiliary populations, nor any difference in the ages of female and male post-hatch or feeding group auxiliaries, in either year.

Male post-hatch group auxiliaries included several individuals who were the social and genetic sons of both breeders (N = 6 in 2001 and 5 in 2002), and also included an individual in his natal territory with his father and his replacement mate, an individual in his natal territory with his social mother and her replacement mate, two different individuals (one for both years) in a natal territory with a half-brother and his mate, a male that had moved with his older brother, and three immigrants unrelated to both breeders. All 20 (total in the two years) female post-hatch auxiliaries were social and/or known genetic daughters of female breeders, and 18 were also social daughters of male breeders (the two exceptions were a single female [in both years] at home with her mother and a replacement for her father [FX; Appendix A 2b in Caffrey and Peterson 2015]).

Feeding groups differed from post-hatch groups in that in 2001, two male auxiliaries (in different groups, a social son of both breeders and an immigrant; Table 1) were never observed to make a nestling feeding trip; in 2002, neither were two females (in different groups, both social daughters of both breeders; Table 1). There was little difference in the genetic relationships to breeders of post-hatch and feeding group auxiliaries (Fig. 2).

**Fig. 2.**
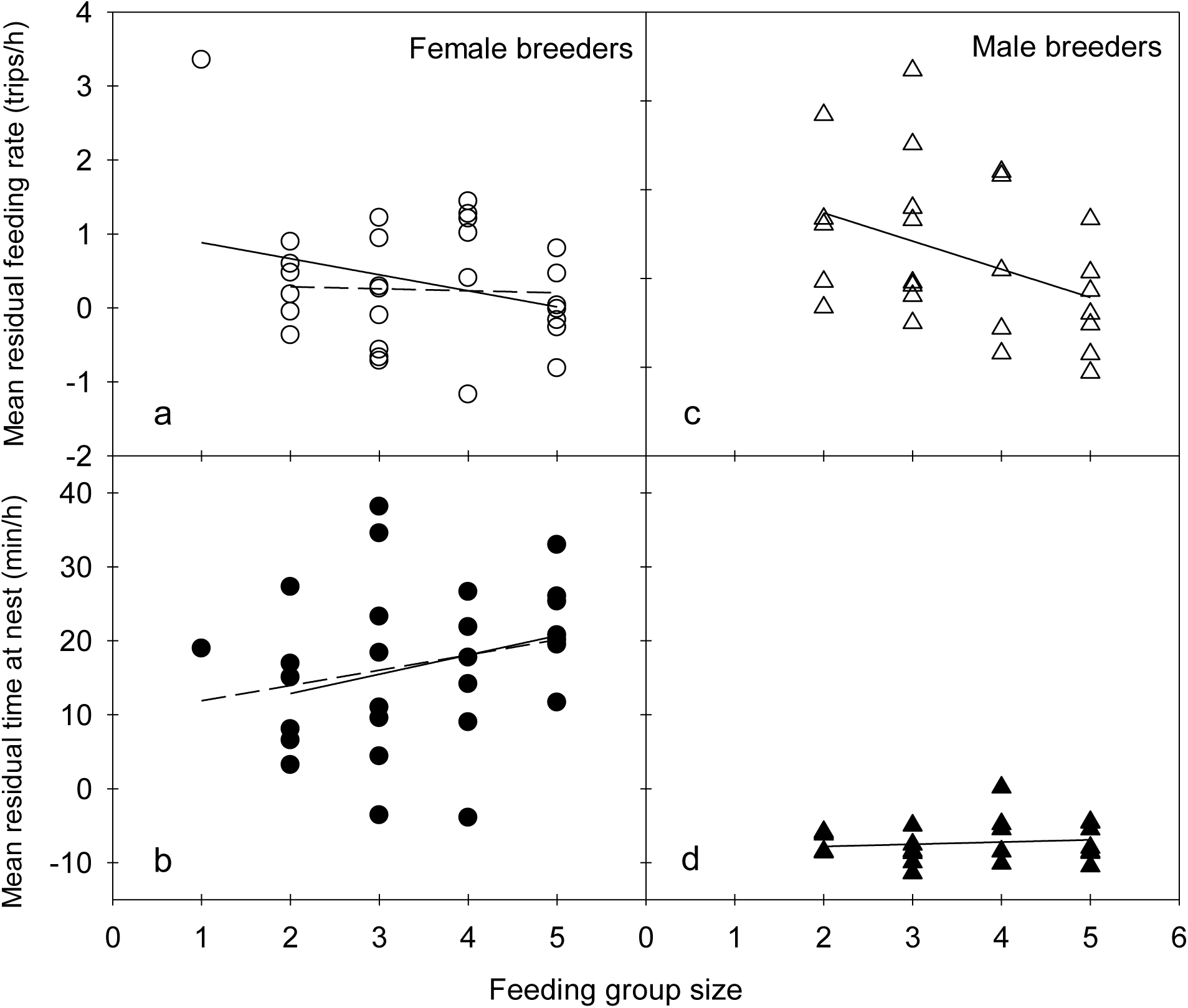
Mean (±95%CI) genetic relatedness to breeders of auxiliary members of pre-hatch groups, pre-hatch auxiliaries that left groups, and members of post-hatch and feeding groups. Solid bars depict relatedness to female breeders; unfilled bars that to male breeders. Numerals indicate sample sizes. Pre-hatch, dispersal, and post-hatch data previously reported (Caffrey and Peterson 2015).

Post-hatch and feeding group sizes changed in four groups upon the disappearance or death of group members (Table 1): In 2001, the male breeder in one group of 3 disappeared six days after hatching had begun in his nest, a 3-year old female auxiliary of one group disappeared 25 days post-hatching, and in another group, the one-year old female auxiliary and male breeder were killed, in separate incidents, after about two weeks of nestling feeding (Appendix A 1a [Group 29] in Caffrey and Peterson 2015). In 2002, a male breeder disappeared after five days of nestling feeding.

### Hatching and Nestling Periods

The hatching of eggs was signaled by infrequent nestling-feeding trips, changes in the behavior of female breeders upon returning after brief breaks (breaks not involving foraging for nestlings; Appendix 1a), and the behavior of some auxiliaries at some nests (Appendix 1b). Activity at most nests continued at relatively low levels for the first few days thereafter, with females sitting for long periods of time (Fig. 3). We saw two visits to nests on the first day of hatching by extra-group individuals (Appendix 1c).

**Fig. 3.**
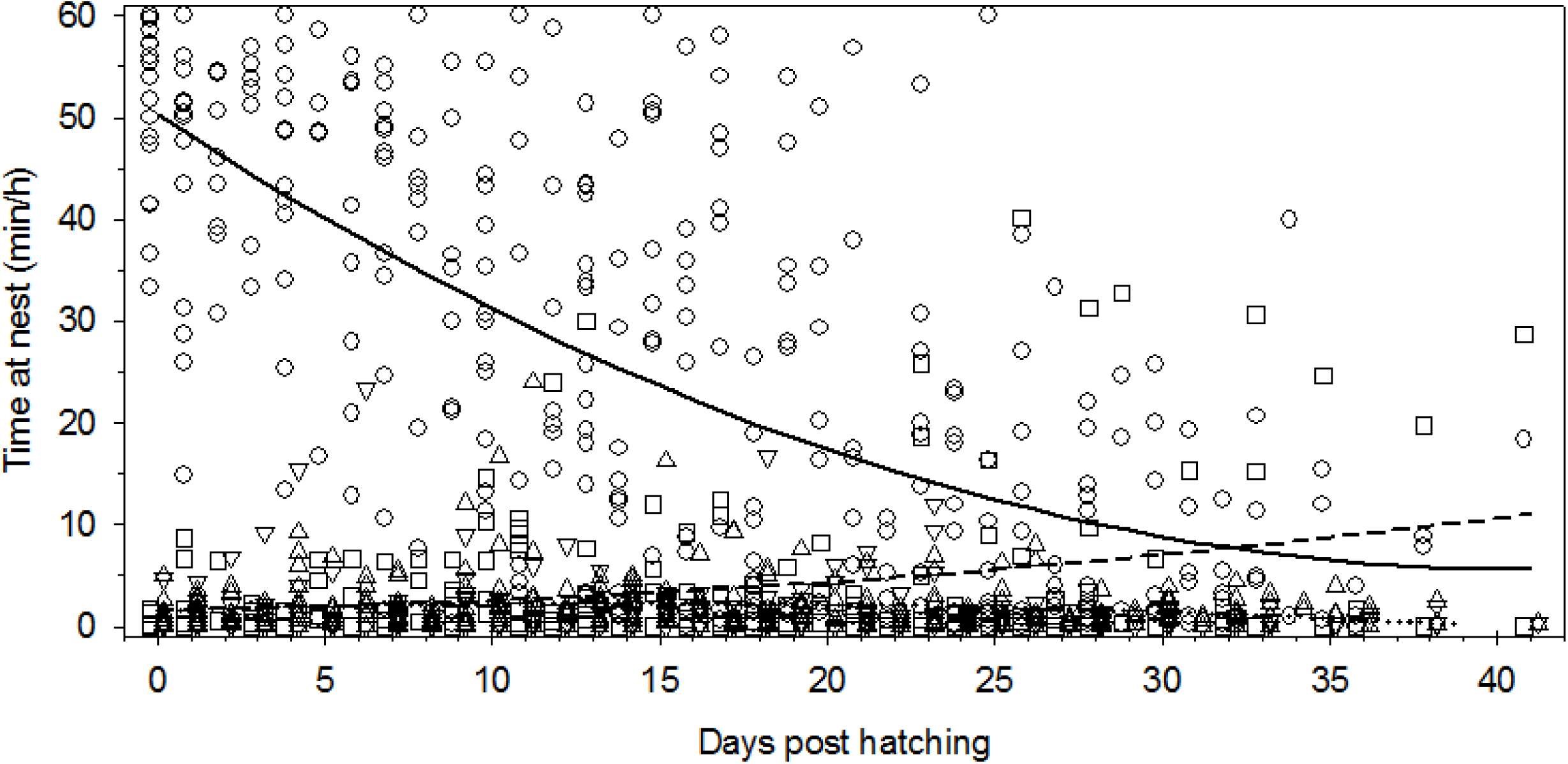
Time spent at and in nests during nestling stages by breeding and auxiliary crows; sex and status symbols as in Fig. 1. Solid curve is best-fit quadratic for female breeders; dashed is that for female auxiliaries.

Nestling periods (first day of hatching to last day of fledging) ranged from 29-42 days, with means of 36.7 days (N = 7) in 2001 and 34.2 days (N = 8) in 2002.

### Nestling Care

Because we analyzed nestling feeding and time-at-nest data for only one attempt per group per year, the total number of auxiliaries included in our feeding group data set was 37 (Table 1; *cf* 40 in Table 2 of Caffrey and Peterson 2015). Across group members, feeding contributions were highly variable. Four auxiliaries made no feeding trips during nest watches, and several made few (including two male auxiliaries, ages 1 and 4, who made only one and two trips, in 881 and 971 min of observation, respectively). Four extra-group individuals were also observed to make nestling-feeding trips (Appendix 2).

Breeders and auxiliaries arrived at nests singly, or with – or at the same time as – another individual, to deliver food to nestlings. During the first 10±2 days after hatching, when female breeders were often in nests (Fig. 3), many nestling feeding trips involved the transfer to female breeders of all or some of the food brought by deliverers (male breeders: proportion of 794 total feeding trips involving handoffs = 15%, of 296 trips in first 10 days = 29%; female auxiliaries (of 320 and 122 trips) = 15% and 28%, respectively, and male auxiliaries (of 336 and 96 trips) = 10% and 23%, respectively). If deliverers had kept some of the food, both they and female breeders then fed nestlings. Breeders arriving at nests with food sometimes passed some to other group members (at nests) who then fed simultaneously, and sometimes it appeared a kind of competitive, party atmosphere at nests as individuals (auxiliaries, mostly) arriving simultaneously with food would jostle with each other to get to nestlings.

In addition to expanded esophageal pouches, we saw several types of identifiable items carried to nests (Appendix 3a). One female breeder cached some type of food item in the exterior of her nest from which she took bits to add to food she was already carrying (to nestlings) over a few days. On several occasions, we saw deliverers of food go right past larger begging nestlings up front to feed smaller ones behind, and on one occasion, a male breeder walk all the way around the nest rim to feed all four (large, begging) nestlings.

Feeding group members, especially female breeders, continued to perform a behavior begun in the later stages of incubation (stand in nest and seemingly hammer repeatedly at spots in cup bottoms; Caffrey et al. 2016a, Results, Incubation); this behavior decreased in frequency as nestlings aged. All feeding group members ate or removed fecal sacs when produced in their presence.

### Time at Nests

Feeding group members spent varying amounts of time on nest rims (when nestlings were small, auxiliaries sometimes also awkwardly got *in* nests). When on rims, crows looked around and looked at and preened nestlings (when nestlings were small, nest visitors would lean down into nests and do something we could not see but that observers independently described as “gentle;” presumably they were preening nestlings), and sometimes preened themselves or other group members.

### Relationships with Measured Variables

We examined potential correlates and determinates of feeding rates and time spent at nests with use of ANOVA, ANCOVA, and multiple regression. Despite the extensive variation, at both individual and nest levels of analysis (Fig. 1), five patterns emerged; two associated with breeder feeding behavior and three with time spent at nests. A sixth finding – no patterns associated with measured individual characteristics – was also clear.

The first pattern, according to a two-way ANOVA of time-adjusted (residual) feeding rates, was that breeders fed nestlings more frequently than auxiliaries (F_1, 88_=15.2, P=0.002; sexes did not differ: main effect F_1, 88_=0.32, P=0.57, sex x status interaction F_1, 88_=0.008, P=0.93; Fig. 1a). The second was that when female and male breeders were considered separately, there were significant trends for breeders of both sexes to reduce their nestling feeding rates when in larger feeding groups (females, *r*=-0.36, P=0.06; males, *r*= -0.34, P=0.09: Fig. 4a and c). (Analogously, a female breeder [AM] increased her rate by a factor of 3.3 when rendered the sole member of a group after 16 days of nestling feeding [Fig. 1a and Appendix 3b].)

**Fig. 4.**
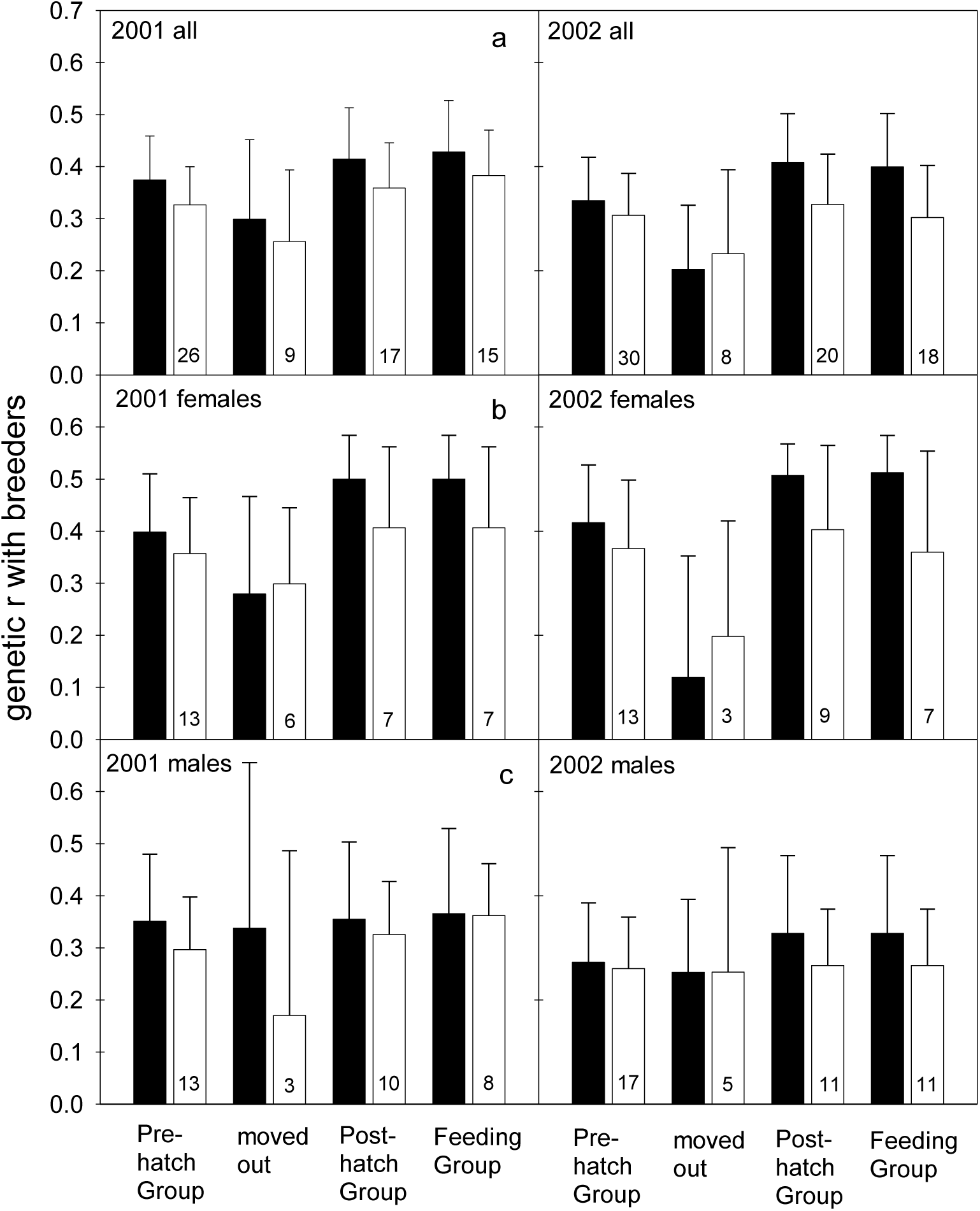
Feeding rates (a and c) and time spent at nests (b and d) of and by breeding crows in feeding groups of different sizes. (a) Correlation with AM (the group of one), *r*= -0.36, P= 0.06 (solid line); without AM, *r*= -0.14, P= 0.49. (b) Correlation with AM, *r*= 0.31, P= 0.11; without AM, *r*= 0.35, P= 0.07 (solid line). (c) *r*= -0.34, P= 0.09.

The third pattern to emerge was that female breeders spent much more time at nests during the nestling period than did male breeders or other group members (Fig. 3); two-way ANOVA results for time spent at the nest during the nestling period were dominated by a strong sex x status interaction (F_1, 88_ = 61.5, P < 0.0001). Female breeders in larger groups spent more time at nests than did female breeders in smaller groups (the fourth pattern: Fig. 4b), and fifth, the time spent at nests by female auxiliaries increased as nestlings got older, despite no change in auxiliary feeding rates (Fig. 3; *r*=0.291; P=0.0001).

Nest-wise, the proportion of time watched during which group members were present at nests was correlated with both post-hatch and feeding group sizes, such that nests of larger groups were attended for more time (*r*=0.405 and 0.508, P=0.050 and 0.011, respectively).

We detected no effects of sex, nesting experience, nest attempt (a, b, or c), post-hatch group size, feeding group size, total brood mass at marking, mean nestling condition at marking, the number of fledglings, or average relatedness to nestlings on breeder nestling-feeding rates (all P>0.15). Feeding rates of auxiliaries were not related to any variable tested, including sex, age, nesting attempt, post-hatch group size, feeding group size, total brood mass at marking, mean nestling condition at marking, number of fledglings, social or genetic relatedness to breeders, or average relatedness to nestlings (all P>0.15).

Comparisons among nest-wise feeding rates (Fig. 1b) did not demonstrate any effects of hatch date, breeder nesting experience, post-hatch group size, feeding group size, total brood mass at marking, mean nestling condition, length of nestling period, or number fledged (all P>0.15).

### Responses to Disappearances

For groups as units, there was a trend for feeding rates to decline after feeding group members disappeared, but variation was high (Fig. 5a); the result of the variable behavior of individuals (Fig. 5b-d and Appendix 3c-e).

**Fig. 5.**
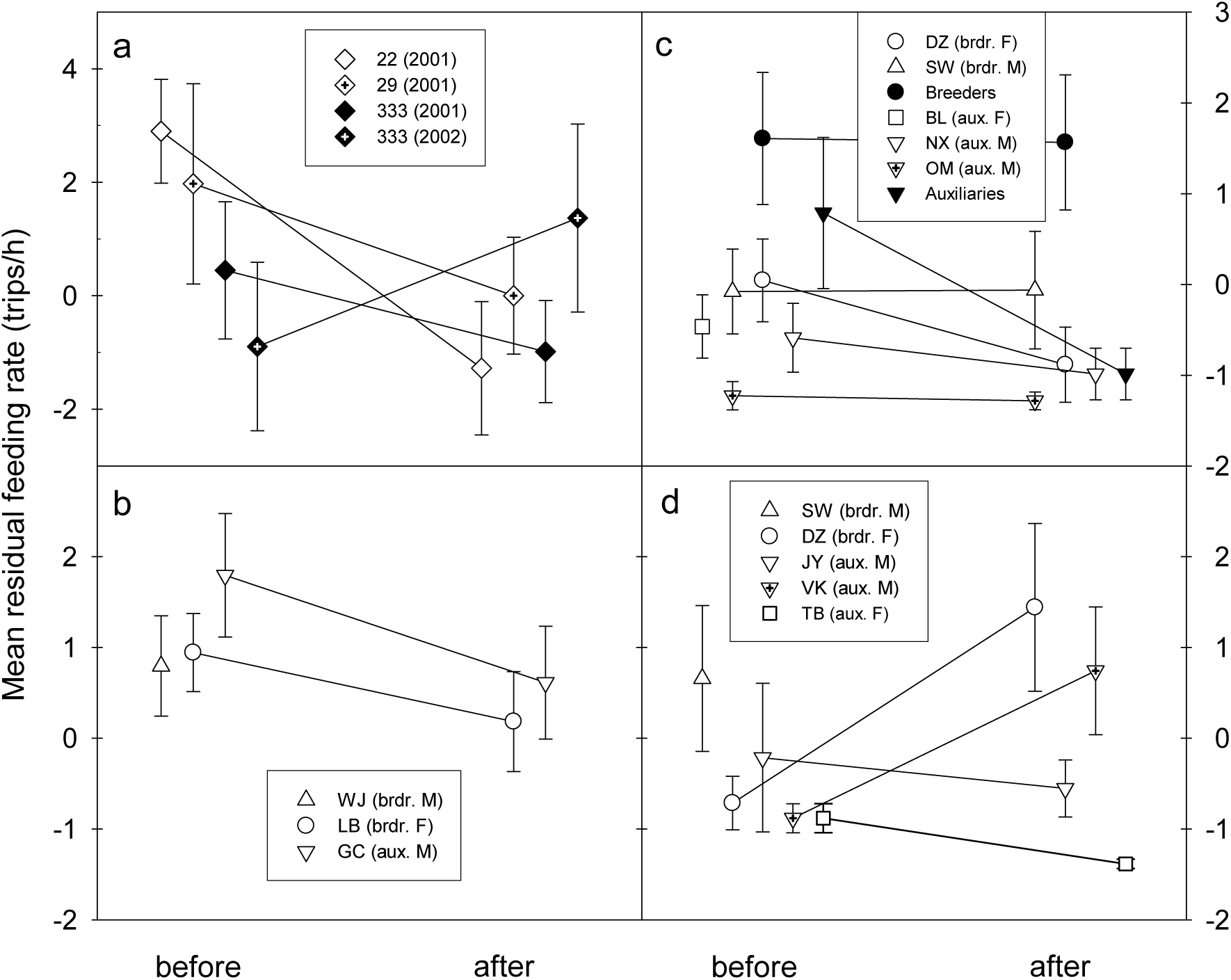
Changes in feeding rates of group members subsequent to the disappearance of others. (a) Nest-wise. (b) Group 22 in 2001. (c) Group 333 in 2001. (d) Group 333 in 2002.

## DISCUSSION

### Post-hatch and Feeding Groups

Because of the dispersal out of groups during the week or so preceding hatching by some auxiliaries, post-hatch crow groups were slightly smaller in size and somewhat less diverse than pre-hatch groups (Table 1 in Caffrey and Peterson 2015), yet the composition of post-hatch and feeding groups was still highly variable (Table 1). Much of the variety stemmed from the diversity in composition of the male post-hatch auxiliary population, which included social and genetic sons of breeders, stepsons, siblings, half-siblings, and immigrants. In striking contrast, it is likely that all post-hatch female auxiliaries were the genetic daughters of female breeders, and many also the genetic daughters of male breeders. That there was no sex bias or differences in the ages of female and male auxiliaries contrasts with the auxiliary population in Ithaca, NY, where older males outnumber older females (Townsend et al. 2009).

Two male and two female post-hatch auxiliaries, in 2001 and 2002 respectively, chose to not contribute to nestling feeding (at least during nest watches), slightly increasing and decreasing, respectively and relative to that of post-hatch auxiliaries, average relatedness of feeding group members to breeders, but not in significant ways (Fig. 2).

### Nestling Care, Patterns

Hatching is asynchronous in American Crows (Verbeek and Caffrey 2002), and so nestling food demand begins low and increases over days as nestlings hatch and grow. The pattern by which nest-level feeding rate changed over time in Stillwater, OK (Fig. 1b) was nearly identical to the pattern observed in a population of Western American Crows in Los Angeles, CA (Caffrey 1999).

### Breeder Decisions and Compensation

Within and across the two years, breeders in our population fed nestlings significantly more times per hour than did auxiliaries; a common pattern among cooperative breeders, and predicted by current theory (Brouwer et al. 2014). Although females and males within many pairs split their feeding effort relatively evenly, the contributions of individual breeders to nestling feeding varied widely: for males, from 1-70% of all observed feeding trips; for females, 12-68% (Appendix 3f).

When female and male breeders were considered separately, their feeding rates were unrelated to any measured variable or any measure of productivity (below), yet a pattern of adjustment to group size was clear: many breeders of both sexes reduced their feeding rates when assisted by additional feeders (Fig. 4a and c). Breeders in a population of Western American Crows responded in the same way to assistance in nestling feeding (Caffrey 1999), whereas Carrion Crow breeders apparently choose to carry less food per trip (Canestrari et al. 2007). These “load-lightening” responses (Crick 1992) make sense in light of Canestrari et al. (2007)’s demonstration that Carrion Crow breeders (and helpers) lose weight over nestling periods in proportion to their calculated nestling-feeding “workloads;” mean loss over nestling period = 5.4% of body weight.

Theory predicts that, on average, breeders will compensate for care by helpers – they will reduce their own feeding effort – when brood reduction via starvation is rare and group size is not positively related to nesting success (often measured as the number of young fledged; Hatchwell 1999). In other words, breeders with assistance can - and do - choose to lighten their own loads when additional food to nestlings is unlikely to increase fledging success (Cockburn 1998, Hatchwell 1999). We have no direct evidence regarding the degree to which brood reduction occurred or influenced fledging success in our study population, but it is an important source of Carrion Crow chick mortality (Canestrari et al. 2007), and its occurrence among American Crow nests in Ithaca, NY, is “rampant” (KJ McGowan, pers. comm.). Clutch size for American Crows is usually 4-5 (mean = 4.7), of which about four usually hatch (McGowan 2001), and the mean numbers of young fledged from nests in our population in 2001 and 2002 were 2.09±1.58 (sd; n=11 nests) and 3.08±1.19 (n=13), respectively; apparently brood reduction *did* constrain fledgling production in Stillwater. Post-hatch or feeding group sizes did not influence fledging success in our population in 2001 or 2002 (a mere snapshot in the lives of these birds), and so it is possible that breeders with help feeding nestlings might have fledged more young had they fed at the higher rates they would have without help. This suggests that the long lives of these birds made the banking of reserves for the future more profitable, on average, than investing in additional current offspring. We wonder, too, if considerations having to do with incipient and future group size may have also influenced breeder effort decisions (i.e., are there consequential costs to too many fledglings?).

Compensating breeders in other species have been shown to benefit in ways unlikely to have influenced the nestling-feeding decisions of breeders in our population. Pairs did not ever attempt to produce more than one brood of fledglings per year, and so enhanced multiple-brooding opportunities were presumably unimportant, as were considerations having to do with the success of renesting attempts: of 30 groups over five years whose a-attempt nests failed (of 83 total groups; eight failed during nest building, 17 during incubation, and five post-hatching), the presence of auxiliaries was not related to the likelihood of renesting or the success of renesting attempts (CC, unpubl. data). (That 21 of 27 renesting attempts, including all three of those having failed post-hatching, succeeded in fledging young [CC, unpubl. data] contrasts with the complete lack of success of renesting attempts in a population of Western American Crows; Caffrey 2000a.) The lack of an effect of auxiliaries on renesting patterns contrasts with the findings for Carrion Crows, in which group size is positively correlated with the probability of renesting after failure (Canestrari et al. 2008). Prior to the arrival in our population of West Nile virus in late summer of 2002 (Caffrey et al. 2005), annual survivorship of breeders had been high: for the four years 1998 through 2001, annual survivorship ranged from 88-100% for both sexes and was equal for both overall: 53 of 56 females (95%) and 53 of 56 males survived to the following year (the one male whose fate was known had been hit by a car; Appendix A 1a in Caffrey and Peterson 2015), and so it is also unlikely that increased survivorship, *per se,* was the benefit being sought by compensating breeders, as has been described for other species (Hatchwell 1999, Dickinson and Hatchwell 2004). Rather, given the luxury of being able to do so, these long-lived, iteroparous organisms were likely saving up for greater investment in future reproductive attempts, an idea also suggested by Canestrari et al. (2007, 2010), and supported by the fact that AM – the female left on her own with nestlings 2.5 weeks old - had not been working at maximum capacity prior to the disappearance of her two other group members (Fig. 1a and Appendix 3b; also the case for breeders in a population of Western American Crows [Caffrey 1999]). AM’s response to being provided with food – to invest in self-maintenance rather than increase the rate to which she had pushed herself (Fig. 1a and Appendix 3b) – mirrors the responses of Carrion Crows (breeders and helpers used the extra energy from food supplementation to reduce, by significant amounts, the weight they would have otherwise lost via feeding nestlings [Canestrari et al. 2007]), and supports, too, the hypothesis that breeders were managing annual energy budgets so as to maximize opportunities over the long term.

In light of Brouwer et al. (2014)’s finding that Red-winged Fairy-wren (*Malurus elegans*) breeders compensate as a function of the number of male helpers but not female, we looked for but found no relationships between breeder feeding rates and auxiliary sex ratio, the number of female auxiliaries in feeding groups (range = 0-2), or the number of male auxiliaries (range = 0-2).

### Time at Nest

That female breeders spent significantly more time in and at nests than other group members during nestling periods (Fig. 3) was not surprising, as females spent much of their time in the first 7-14 days post-hatching brooding nestlings (or sometimes, with wings spread and drooped, shading nestlings or protecting them from rain). Thereafter they spent more time than others preening and otherwise interacting with nestlings (including actively keeping large, flapping nestlings from leaving nests, sometimes using their feet), or (seemingly) simply hanging out at nests. This appears to be one of the roles of female breeders during nesting, as we found a significant relationship in the general patterns of time spent at active nests for the 19 individuals for which we had data for both years (Fig. 6a).

**Fig. 6.**
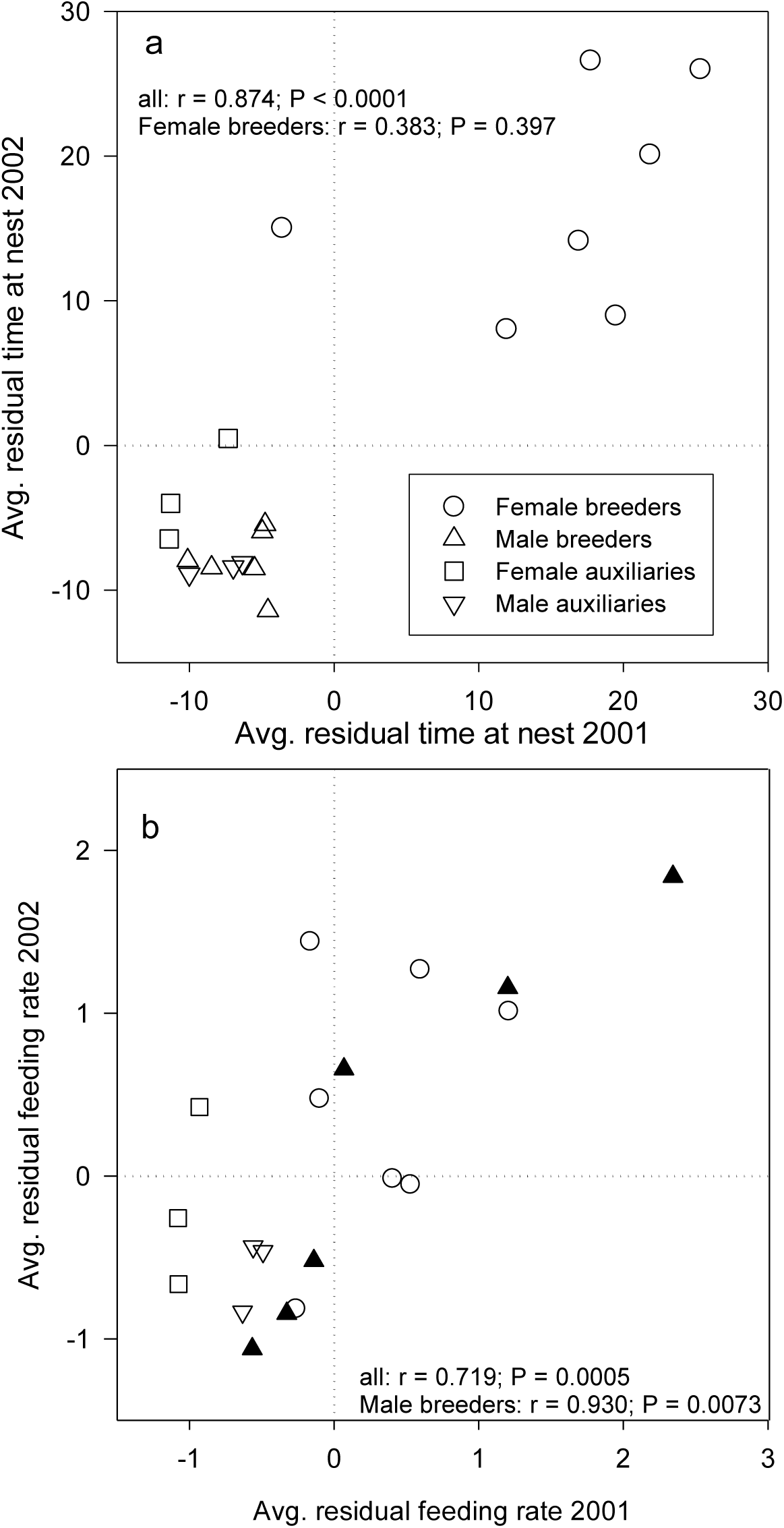
Relationships between behavior in 2001 and 2002 for feeding group members observed in both years. (a) Time spent at nests. (b) Nestling feeding rates.

Considerations having to do with time spent at nests by female breeders are myriad and not straightforward in their interpretation. Female breeders in larger feeding groups spent more time at and in nests, on average, than did females in smaller groups (Fig. 4b). (With the inclusion of AM – again, the female rendered a group of one upon the deaths of groupmates [Appendix 3b, and Appendix A 1a in Caffrey and Peterson 2015] – the correlation was not significant, but as her time-at-nest was confounded by the increased time she spent feeding nestlings, a better index of decisions by female breeders regarding the time they spent at nests as functions of group size is the correlation excluding AM [Fig. 4b; *r*=0.35, P=0.07].) Because female breeder time-at-nest was tightly correlated with nest-wise time-at-nest (*r*=0.929, P<0.0001), the total percent of time nestlings were attended was also a function of feeding group size (and because of the tight correlation between feeding group and post-hatch group sizes, female breeder and nest-level time-at-nest were both also positively correlated with post-hatch group size).

As female breeders in larger groups fed nestlings at lower rates than did those in smaller groups, on average (Fig. 4a), the increased time they spent at nests was *not* related to feeding nestlings; rather, female breeders in larger groups spent more time with nestlings during the first 10 days post-hatching, the time nestlings are most vulnerable to external influences (first 10 days, *r*=0.336, P=0.003; thereafter, *r*=0.297, P=0.141). (As above, it was during the first 10 days post-hatching that most handoffs of nestling food to female breeders occurred, and the behavior of females upon acceptance [the ways in which they stood up and positioned themselves] suggested that nestling thermoregulation was important.) Yet, greater amounts of time spent at nests by female breeders, or other group members, were not positively related to any measure of productivity, and were in fact significantly *negatively* correlated with the mean condition of nestlings (female breeders, *r*=-0.720, P=0.006; total, *r*=-0.805, P=0.0009) and the number of young fledged (female breeders, *r*=-0.397, P=0.055; total, *r*=-0.424, P=0.039). The negative relationships likely reflected extended time spent with some last (weakest) nestlings to fledge in 2001 and 2002 (the lengths of nestling periods were negatively correlated with mean nestling condition [*r*=-0.64, N=10, P=0.046], and number of young fledged [*r*=-0.37, N=15, P=0.180]), and so the extra time female breeders in larger groups spent at nests did not manifest in enhanced current fledgling production.

Individual auxiliaries of both sexes varied in their seeming interest in nestlings; some spent little to no time at nests other than that involved in the transfer of food while others spent many minutes (one more than 36; she was there when we arrived) at nests, gazing down into them, preening nestlings, and just hanging out. Female auxiliaries tended to spend more time at nests than did male auxiliaries (with a few exceptions), likely in keeping with their future roles as female breeders (above: Fig. 6a). The significant increase in the time female auxiliaries spent at nests as nestlings aged (Fig. 3) was driven mostly by a single individual – a two-year old social daughter of both breeders – who spent large amounts of time on the nest rim, looking at nestlings, looking around, preening nestlings, preening herself, and seemingly just passing time (two other female auxiliaries were also notable in their interest in, and time spent preening, nestlings).

### Nestling Care; Individual Decisions and Proximate and Ultimate Considerations

As eggs hatched in nests of crows in Stillwater, breeders and auxiliaries who chose to stay for nestling periods faced their next sets of decisions, some having to do with direct nestling care: To feed or not? If so, on what kind of schedule? How much energy to commit to nestling care this season? The high level of diversity in the strategies we observed suggests the proximate and ultimate mechanisms at work must have also been diverse. Our data shed light on but a few.

On the proximate side and broadly speaking, a significant proportion of the variation in individual contributions to nestling care was attributable to the different roles played by different group members during nesting seasons; despite small sample sizes, we found that feeding group members of the same sex and status (breeder or auxiliary) behaved in the same general ways at nests during nestling stages in the two years (Fig. 6a and b). (The aberrant female breeder [below zero in 2001] in Fig. 6a was paired with a male described in feeding watch notes that year as “A <u>great</u> father and mate!” He is the topmost male in Fig. 6b; his extremely high level of nestling care was likely related to her reduced presence at their nest in 2001, although we do not know the causality.) With regard to feeding rates, we were unable to test for differences within sex for auxiliaries, but individual breeders were consistent in their behavior across years (Fig. 6b); for males, significantly so. The female plotted at the top left of Fig. 6b is DZ, whose mate disappeared after five days of nestling feeding in 2002; her feeding rate increased significantly thereafter (Fig. 5d and below; Responses to Disappearances). Without DZ, the correlation between nestling feeding rates in the two years for female breeders was *r*= 0.676 (P>0.1). The nestling feeding behavior of the male at the lower left of Figure 6b (UB; Appendix A 2g in Caffrey and Peterson 2015) was possibly influenced by his suffering from a recurrent leg problem during the nesting seasons of 2001 and 2002; Carrion Crow breeders also respond to physical handicap by reducing nestling care (Baglione et al. 2010). The repeatable variation among breeders, especially males, is strongly suggestive of different “personalities” (Dall et al. 2004, Réale et al. 2007). We did not measure crow behavior to test for personality traits, yet we knew them to “exhibit between-individual differences” (Carter et al. 2013), and the consistency in breeder care of nestlings across years would appear to meet the criteria of the definition: inter-individual differences in behavior being maintained over time (Réale et al. 2007, Carere and Locurto 2011, Webster and Ward 2011, Wolf and Weissing 2012, Carter et al. 2013); in this case, a “behavior” as complex as care of nestlings, and a time periods of years.

Personalities have been reported in a wide range of animals (Réale et al. 2007, Bergmüller et al. 2010, Carere and Locurto 2011 and references therein, Wolf and Weissing 2012 and references therein), and they were manifest in the crows in our population at two levels: group and individual. Although within-group variation existed in individual behavior, some groups differed markedly from others in reluctance, or not, to approach or return to bait, for example. In addition, although virtually all population members had experienced us under similar trapping and nestling-marking conditions, thereafter members of some groups seemed indifferent to our presence, members of other groups did not like and were extremely wary of us, and members of others not only acknowledged us but sometimes approached our vehicles and accepted peanuts tossed from windows. (As such, group members were likely “conforming” with each other; Webster and Ward 2011.) Such “group personalities” (Koski and Burkart 2015) have been described for Common Marmosets (*Callithrix jacchus*; Koski and Burkart 2015, chimpanzees (*Pan troglodytes*; Cronin et al. 2014), and Common Ravens (*Corvus corax*; Miller et al. 2016); we wonder if the lack of such a finding for Carrion Crow/Hooded Crow hybrids (*C. corone corone/C. C. cornix*) might have been related to the artificiality of the materials used in the tests (Miller et al. 2016). No individual crows in our population behaved the same all the time, yet some individuals were notably more bold and aggressive than others, some more inquisitive than others, some more playful, some seemed more prone to routine, some (mostly females) were especially gentle and caring with nestlings, and a few seemed particularly smart, e.g., the individual that made a tool with which to probe a space too small for its head (Caffrey 2000b), and AM (Appendix 4, and 3b, and Appendix A 1a in Caffrey and Peterson 2015).

### Breeders

Across all breeders, the rate at which individuals fed nestlings varied considerably and was not influenced by their sex, their experience with current mates, whether nestlings were in first- or later-attempt nests, post-hatch group size, feeding group size, numbers and ratios of auxiliary sexes, their average relatedness to the nestlings in their nests, the average body condition of the nestlings in their nests, the total mass of their brood at marking, or the number of their nestlings that fledged. We found no evidence for current reproductive benefits associated with having extra individuals feeding or otherwise attending to the care of their social nestlings. Instead, as above, when assisted in nestling feeding by auxiliaries, many breeders of both sexes exploited the opportunity to maximize future reproductive benefits: they chose to reduce their feeding rates rather than use auxiliary feeding contributions to provide current nestlings with additional food (that males may have compensated moreso than females may reflect their greater uncertainty of parentage [Heinsohn 2004]). Thus, added to the potential net benefits to breeders to allowing offspring to remain home for up to several years (Caffrey and Peterson 2015), were the future benefits associated with exploiting their contributions, and those of others, and reducing current nestling-feeding costs.

Nest-level (total) feeding rates were also unrelated to any variable measured, including total brood mass at marking and the number of young fledged per nest; an unexpected result given that the total energy demand in nests of larger brood sizes *had* to have been higher than the demand of smaller broods. If the begging intensity of nestlings was a function of their hunger (which is known to be the case for at least some other birds [Wright 1998], including another crow [Carrion; Canestrari et al. 2010]), and breeder feeding effort was being influenced by nestling begging intensity (which is known to be the case for at least some other birds [Wright 1998, Wright et al. 2010, Young et al. 2013], including another crow [Carrion; Canestrari et al. 2010]), then it *had* to be the case that crows in groups with larger brood sizes were, on average, bringing more food per feeding trip than were crows in groups with smaller brood sizes (we found no correlation between brood size at marking and total brood mass or mean nestling body condition). Nestlings in physical contact with other nestlings may have had reduced thermoregulatory costs and so nestlings in larger broods may have grown more efficiently, yet reduced maintenance costs were unlikely to have offset the greater energetic demand of increased numbers of growing bodies. With the use of cameras and the consequent ability to observe fine details of nestling feeding, Canestrari et al. (2005, 2007, 2010) found that Carrion Crows indeed carry different amounts of food, and so feed different numbers of nestlings, on different feeding trips. We suspect the feeders of larger broods in our population were, on average, carrying more food per trip than feeders of smaller ones.

Brouwer et al. (2014) found that Fairy-wrens behave differently in response to the sex makeup of nestling co-feeders: all members reduce their contributions, on average, with increasing numbers of males, but not so with females. Nestlings were therefore fed at higher rates in the presence of female helpers (Brouwer et al. 2014). We looked for such an effect but found suggestive support for the opposite: the number of male auxiliary nestling feeders was weakly positively correlated with total (nest-wise) feeding rate (*r*=0.463, P=0.023).

### Auxiliaries

One feature of the auxiliary crow population in Stillwater, OK (and presumably all crow populations) not often included in considerations of helper costs and benefits is that yearlings are not reproductively mature. As such, definitions of delayed dispersal such as ‘remaining in natal areas past being competent to reproduce’ (e.g., in Ekman et al. 2004) are not even applicable. We knew little about the whereabouts or behavior of individuals that dispersed out of groups (and out of our population) for short or extended periods, yet group living appeared to be an integral part of the lives of our crows and yearlings seemingly had little choice but to be members of others’ groups. (Of w*hose* groups they chose to be members was a separate issue; Caffrey and Peterson 2015.)

For post-hatch auxiliary crows, potential routes to current and future fitness benefits were demonstrable and wide-ranging (Caffrey and Peterson 2015), and so auxiliaries stood to gain by way of being auxiliaries. Not so easily identifiable are the possible factors underlying the huge range of “investments” (Bergmüller et al. 2007) made to nestling care by post-hatch auxiliaries: Some fed at rates equal to or higher than those of breeders; extreme highs included 56% of 54 trips (an adult daughter of both breeders), 47% of 81 trips (an adult social daughter of both breeders [the genetic daughter of the female but not the male]; she made only 6% of 64 the previous year, as a yearling), 45% of 51 trips (an adult son of both breeders; below, Responses to Disappearances [GC] and Appendix 3c), and 41% of 140 trips (an unmarked yearling; likely the social offspring of the breeders). Other auxiliaries made few feeding trips and some may have made none at all (a situation similar to that in Carrion Crows; Baglione et al. 2010); extreme lows included, for three auxiliaries (a yearling son of both breeders, and two daughters [an adult and a yearling] of both breeders), a single feeding trip of 123 (0.8%), 33 (3%), and 33 (3%), respectively, and for a younger brother of the male breeder (an adult male: KR; Appendix A 2a in Caffrey and Peterson 2015), two of 64 (3%). Although possible explanations for the evolutionary maintenance of individual variation in behavior have been proposed (e.g., Dall et al. 2004, Sih et al. 2004, Réale et al. 2007, Taborsky and Oliveira 2012), including variation in cooperative behavior (e.g., Komdeur 2006, Bergmüller et al. 2010, Wolf and Weissing 2012, Dingemanse and Araya-Ajoy 2015), we know of no case made for the degree of within-population diversity that we observed in the nestling-care behavior of auxiliary crows.

As above, the rate at which auxiliaries fed nestlings was not significantly influenced by any measured variable. Feeding group members did not, on average, enhance the production of nondescendant kin (at least in terms of the number of young fledged per nest) via their presence or feeding contributions in 2001 or 2002, and so increasing (current) indirect benefits did not appear to be the objective of their behavior (although for auxiliaries assisting genetic parents who chose to then compensate, such benefits would theoretically accrue in the future).

If auxiliaries had to “pay” to stay in groups (Gaston 1978, Kokko et al. 2002), the currency was unlikely to have been nestling feeding, as we found no evidence of cheating (false feeding, i.e., arriving at nests with food but consuming it rather than delivering it to begging nestlings; Young et al. 2013), as would be expected if auxiliaries were to be perceived as meeting some minimum level of investment, or as investing more than others (Young et al. 2013), and seen in Carrion Crows (Canestrari et al. 2010). Ninety percent of 852 trips to nests by male breeders in our population involved the feeding of nestlings, and the majority of the balance were trips wherein food was delivered to and eaten by brooding females (not passed to nestlings). Nestling-feeding trips accounted for 86% of all trips by male auxiliaries (n=381) and 83% of those by female auxiliaries (n=372); again some of the balance included feeding brooding females, and some non-feeding trips appeared to be quick “checks,” but some individual auxiliaries, especially females, regularly visited nests without food, to attend to nestlings, interact with female breeders, or just hang out (similar to the sex bias to non-feeding trips in Chestnut-crowned Babblers (*Pomatostomus ruficeps*; Young et al. 2013). Of the 1496 trips to nests by female breeders, food was delivered to nestlings on 69% of them, and there was no indication that females had arrived with food and swallowed it on any of the 453 non-feeding trips. However, we once saw a female breeder, after the auxiliary who made the feeding trip had departed, remove some of the food from a nestling’s mouth and eat it herself. We also saw no evidence of punishment or retribution (Cant 2011, Young et al. 2013) toward auxiliaries who contributed little to no nestling care during observation periods, and no evidence of interference among auxiliaries (*sensu* Carlisle and Zahavi 1986). (We *did* see two female breeders repeatedly intervene in the feeding of nestlings by their mates; we include an example in Appendix 3g).

The diversity of nestling-season options pursued by individual post-hatch auxiliaries in our population ran the gamut usually described in among-species comparisons of direct fitness components (e.g., Heinsohn and Legge 1999, Heinsohn 2004), and so analogously, we suggest that the costs and benefits associated with group membership and nestling care varied for different individual crows *within* our population. In other words, the differences we observed among individuals were no accident (*sensu* Hirsch 1963). Feeding nestlings is costly, and crows are smart, long-lived, behaviorally flexible organisms; feeders *should* adjust investments as functions of both current and long-term considerations. Behavior is the outcome of the integration of many of an organism’s biological systems (Hirsch 1963), and individual crows in Stillwater differed in uncountable ways, including: genotype (Hirsch 1963, Komdeur 2006), sex, age (although we found no evidence that yearlings were less capable contributors than adults; references in Kramer and Otarola-Castillo 2015) and weight (Komdeur 2006 and references therein), past experiences and relatedness to breeders and nestlings, and presumably also (as addressed in the literature) body condition (Dufty and Belthoff 2001, Sih et al. 2004), personality or temperament (Quinn et al. 2001, Sih et al. 2004, Dall et al. 2004, Réale et al. 2007, Ensminger and Westneat 2012), circulating hormone levels (Komdeur 2006 and Réale et al. 2007, and references therein), developmental and maternal and/or parental effects (Komdeur 2006 and references therein, Reddon 2011), gregariousness (Nuñez et al. 2014), “state” (the combination of factors determining current behavioral decisions, including stochastic influences [Dall et al. 2004] and individual “time programmes;” Helm et al. 2006), quality (Heinsohn 2004, and see below), “emotional state” (Dall et al. 2004 and references therein), “pace of life” (synergistic suite of behavioral, physiological, and life history traits; Réale et al. 2010), “stakes” in other individuals in their groups (extent to which fitness depends on that of others; Roberts 2005), and “social competence” (ability to optimize behavior in response to social information; Taborsky and Oliveira 2012). The optimal “social strategies” (Helms Cahan et al. 2002) of individuals may have also been in flux.

Differences in the step-wise decision “trajectories” (Helms Cahan et al. 2002) of individual auxiliaries – whether to leave pre-hatch groups or stay, whether to feed nestlings or not, and how much nestling care to actually contribute – may have had something to do with differences in individual “quality” (Barclay and Kern Reeve 2012), but the relationships between our observations and the assumptions and predictions of Barclay and Kern Reeve’s (2012) model are not straightforward. The nestling-feeding contributions of auxiliaries in our population ranged from less than 1% to 56% of group total feeding trips. Theory predicts that individuals will vary in their help as a function of their own quality, the nature of the assistance they are providing, and the nature of the opportunities helpers are forgoing in order to help (Barclay and Kern Reeve 2012): if the help being provided is quality-dependent, such that higher-quality individuals are able to provide it at lower “performance cost” than lower-quality individuals, then all else being equal (including the benefits of helping), higher-quality individuals should help more than lower-quality individuals (implicit in this argument is that they *will* simply because they *can*). Yet time spent helping is time unavailable to spend on other activities, such as foraging or pursuing reproductive opportunities, and because the nature of these lost opportunities differs, such “opportunity costs” likely differ for individuals of different quality (Barclay and Kern Reeve 2012): if the lost opportunity is satiable, for example one’s own foraging requirements, then higher-quality individuals should become sated sooner and thereby pay lower opportunity costs for helping to feed nestlings. If the lost opportunity is one with increasing potential benefits available for increased commitment (e.g., time spent attempting to attract mates), then higher-quality individuals would pay higher opportunity costs and lower-quality individuals would be expected to help more, especially if the help they are providing is not very quality-dependent (and so performance costs are not prohibitive; Barclay and Kern Reeve 2012). We do not think that foraging for nestlings was particularly difficult; a feeding group member with a broken leg (Appendix 3f) and ones burdened with inappropriate patagial tags (Caffrey 2002b) fed nestlings at average rates, feeders sometimes returned to nests after only a few minutes of foraging, all nestling feeders spent considerable amounts of their time throughout days loafing, and again, AM was able to triple her feeding rate upon the disappearance of her two group-mates (at no immediate, measurable cost to her fitness: she survived through at least the next two years). As such, the feeding help being provided was likely not particularly quality-dependent. The costs of lost potential reproductive opportunities do not apply to yearling crows contributing variable amounts of effort, and although adult auxiliaries in the earliest-nesting groups may have had up to a week of overlap between hatching in their own groups and the fertile periods of late-nesting (renesting) females in other groups (who ended up successful in fledging young), most auxiliaries were not sacrificing direct opportunities for reproducing by choosing to feed nestlings. Barclay and Kern Reeve (2012) and others (e.g., Carlisle and Zahavi 1986, Zahavi 1990, Putland 2001) have suggested that industrious helpers may be advertising their quality or displaying their parental ability to potential future mates, and, in fact, one male breeder in this study *had* previously assisted his mate (and her previous mate) in raising nestlings (KB; Appendix A 2h in Caffrey and Peterson 2015). Yet the lack (or low rate) of false feeding among auxiliaries (i.e., no “cheating”) has been used to suggest, for other species, that helping does *not* function in a signaling context (Young et al. 2013). The 10 auxiliaries (of 37) in our population that fed nestlings at the greatest rates in 2001 and 2002 included three yearlings (two females and one unmarked [the latter likely fledged from the nest of breeders and delayed dispersal]), six two-year olds (three females and three males), and a four-year old male immigrant. Other than the immigrant (TM; Appendix A 1a and 2e in Caffrey and Peterson 2015), all others were assisting social parents of the opposite sex. Two of the eight individuals were known to be the genetic offspring of their social parents, but one, a two-year old female, was *not* the daughter of the male she was assisting (she and he, upon the death of his mate [the auxiliary’s mother], both ended up pairing with other crows). It is still possible that opportunities to impress possible future mates may have been influencing the nestling-feeding decisions of some auxiliaries – they may have been signaling to *each other* - yet given the stumbling blocks to “social prestige” operating reliably (Wright 2007), it is likely most auxiliary feeding group members were seeking benefits of other kinds.

### Responses to Disappearances

As discussed by Liebl et al. (2016a), empirical attempts at quantifying carer effects on provisioning rates of nestlings in cooperatively-breeding species are rare, and have taken two routes: experimental carer removal and brood enhancement. The two approaches have their pros and cons (Liebl et al. 2016a and references therein, Liebl et al. 2016b); relevant here is that both are unnatural manipulations of natural systems. The disappearances of crows from our population during the nestling season offered a natural test of the impact of feeding group members on “sustenance received by offspring” (Liebl et al. 2016a). At first glance (Fig. 5a), that feeding rates at three of four nests decreased subsequent to the disappearance of feeding group members would seem to support helpers-as-additive (Liebl et al. 2016a and references therein). But details of the identity of disappearers, the timing of disappearances relative to nestling ages, and within-group dynamics and individuality unveil general support for the idea that breeders exploited additional carers to reduce their own feeding rates rather than to provide additional sustenance to nestlings (Kabadayi et al. [2016] draw attention to the potential pitfalls of generalizations without examination of the details of individual behavior). Variation in the behavior of individual crows also played a large part in group responses to member disappearances.

In Group 29 (Fig. 5a), both the female auxiliary and male breeder “disappeared” within days of each other (Appendix 3b). The female breeder (AM) more than tripled her own effort, and survived thereafter, indicating she had not been working at close to the level of which she was capable when assisted by her mate and daughter. Despite AM’s Herculean effort, the above-average feeding rate her nestlings had experienced was reduced to the average rate of the population (Fig.s 5a and 1). The situation at Nest 22 (Fig.s 5a and 5b) was mysterious: For six days, all three group members fed nestlings at high relative rates. At a time when nestling food demand is otherwise increasing (Fig.1), the decrease in the feeding rates of both remaining group members – the mother (LB) and brother (presumably; GC) of the nestlings – might suggest some sort of disturbance (Cockburn 1988 in Lieble et al. 2016a); we wondered if their reduced trips to their nest reflected fear or perceived risk, but their behavior while at, and away from, the nest appeared normal. We could see four nestlings through the day of a heavy storm (Appendix 3c); the following day, LB and GC made seven visits to the nest in 90 minutes of observation: food was brought on two, but the single, weak nestling could not respond to nudges or gentle prods. The nest was abandoned by the next day.

Group 333 epitomizes our understanding of crow behavior: some things make good sense, and many defy logic. In 2001 (Fig. 5c), an auxiliary (BL) disappeared after 25 days post hatching, a time when feeding rates at nests tend to decrease (Fig. 1). The feeding data were complicated by our inability to distinguish the two breeders at all times (Appendix 3d), but together they maintained their pre-disappearance rate, suggesting they were decompensating and making up for BL’s disappearance (again, during a period of population-wide [including the remaining auxiliaries in Group 333] decrease in nestling feeding rates [Fig. 1]).

In 2002 (Fig. 5d), the male breeder (SW) of Group 333 disappeared after only five days of nestling feeding – when feeding rates were just beginning their increases from low starts (Fig.1) – and so the huge, and significant, increase in his mate (DZ)’s provisioning rate makes sense. At the time of SW’s disappearance, DZ had a yearling daughter (TB) and a yearling son (JY) living with her full time, and two two-year old sons sharing residency with Group 333 and one of its next-territory neighbors (Group 55: previous group members, including the father of one of the two-year olds; Appendix 3e). In the days immediately following SW’s disappearance, the nest-level feeding rate at first decreased until JY increased his own. Eight days after SW disappeared, one of the two-year olds began provisioning nestlings. JY thereafter continued to feed nestlings but at a reduced rate; the mean for the population. Baglione et al. (2010) also found that about half of group members not previously contributing to nestling feeding chose to participate in response to the reduced contributions of (handicapped) others, yet the details of the genetic relationships among members of Group 333 make the situation difficult to understand: Reproductive sharing occurred in this group (Appendix 3e) and VK was the genetic son of DZ but not SW, while TB, JY, and the other two-year old male were the genetic offspring of both. As such, VK was the least closely related to the nestlings of all auxiliaries, and, as above, had been spending a lot of time with his father’s group. VK had not made a single nestling feeding trip to Nest 333 before SW disappeared, and was then not observed to do so for another eight days. Thereafter (perhaps in response to the lack of an adequate response by his half-siblings), VK made 37% of feeding trips observed. All else being equal, the three more closely related auxiliaries, especially the two living with DZ (one of which was already feeding nestlings), should have increased *their* provisioning rates of their siblings. Finally, VK was DZ’s son, and so, presumably, not trying to impress her so as to secure a breeding position in the future.

### Concluding Thoughts

In behavioral studies of cooperative breeding, intraspecific differences have been largely ignored or viewed as raw material around adaptive means, rather than the potential products – themselves - of natural selection, leading to the “neglect of whether, how, and why individuals differ in behavior” (Réale et al. 2007). Hopefully, recent acknowledgment of the importance of incorporating individual variation and personality differences into studies of behavior (e.g., Komdeur 2006, Bergmüller et al. 2010, Webster and Ward 2011, Carter et al. 2013, Dingemanse and Araya-Ajoy 2015, Sih et al. 2015), will convince cooperative breeding researchers to bend toward the more “holistic” approach of psychologists (Carter et al. 2013 and references therein), because “If we take individual differences into account our conclusions and explanations of social behavior may change” (Komdeur 2006).

American Crows are long-lived, brainy, behaviorally flexible and complex, socially-savvy animals with senses of humor (e.g., Caffrey 2001) and that exhibit spontaneous, or unsolicited, prosociality (motivational predisposition to perform acts that benefit others, without reward or enforcement [Burkhart et al. 2009]; interpreted as concern for the welfare of others [Bergmüller et al. 2010]; e.g., Appendix A 4 in Caffrey and Peterson 2015) and don’t seem to play by the rules of other cooperative breeders. Kin selection does not underlie their cooperative choices, as attested by all three manifestations of supportive evidence (Komdeur 2006 and references therein): 1) not all auxiliaries contribute to nestling care, 2) auxiliaries (regardless of genetic relationships with breeders) gain direct benefits from group membership (Caffrey and Peterson 2015), and 3) nonbreeders regularly move among groups and may help to raise the unrelated young of unrelated breeders. Individual crows have different personalities, different food preferences (CC, unpubl. data), different capabilities, different agendas, and different types of relationship with lots of other crows; all these phenotypic differences are themselves the result of different genetic, developmental, parental, and past and current environmental influences. (Mothers [or both parents; Reddon 2011 and references therein] may have even purposefully created diverse personalities and behavior among offspring in natal and others’ groups, as a means of increasing their own reproductive success [e.g., Komdeur 2006 and references therein, Reddon 2011].) Fluctuating environmental pressures are associated with maintaining within-population variation in behavioral strategies (Clark and Ehlinger 1987, Réale et al. 2007, Webster and Ward 2011, Taborsky and Oliveira 2012, Dingemanse and Araya-Ajoy 2014), and the weather and foraging conditions in Stillwater varied widely from season to season and, at least in winter, year to year. (Reddon [2011] suggests variable environmental conditions as a reason for diversifying offspring personalities.) On the social side, interactions with social partners with different personalities and their own agendas impose selective pressures on, and influence the fitness consequences of, the behavioral strategies of individuals (Wilson 1998, Réale et al. 2007, Webster and Ward 2011, Taborsky and Oliveira 2012, Dingemanse and Araya-Ajoy 2015, Koski and Burkart 2015), and the social lives of crows in Stillwater were dynamic and complex (Caffrey and Peterson 2015), and rich with opportunities to interact with others in groups of two to hundreds at a time. Behavior can act as a “pacemaker” for evolution, and the process could conceivably be sped up if individuals are exposed to novel selection pressures (Wolf and Weissing 2012); in populations with individuals that vary in phenotypic strengths, the chance that challenges can be met is increased (Sih et al. 2004, Dall et al. 2004, Taborsky and Oliveira 2012). For many animals, such challenges include the continuous and rapidly-occurring changes wrought by humans (Sih et al. 2004, Dall et al. 2004), and especially for crows, possibly also those *induced* by humans [e.g., hunting pressure; Réale et al. 2010]). That crows are notoriously successful at occupying and exploiting human habitats speaks to their ability to keep up adaptively with us; no doubt a function of their intelligence and variable and flexible behavior.

For the reasons above and also because it makes sense that individual crows would adjust their behavior as functions of their own states (Dall et al. 2004), the behavior of others, and current environmental conditions (and with eyes toward the future), it was not surprising that we found no broad-level relationships between measured components of nestling care and phenotypic classes of crows. Unfortunately, the contributory proximate mechanisms and ultimate fitness consequences of particular strategies pursued by individual crows in this study will forever remain unknown, as tragedy rendered those strategies moot: In its spread west across North America in the early 2000s (Hayes et al. 2005), West Nile virus (a non-native invasive pathogen) reached Oklahoma by late summer 2002, and by the end of that year had killed approximately one-third of our population (Caffrey et al. 2005). One year later, most of the crows in Stillwater were dead (Caffrey et al. 2005), and the social and ecological environments in which survivors were left to carry on their lives had been drastically changed.

## ACKNOWLEDGMENTS

We are grateful to all of the people and organizations we acknowledge in Caffrey and Peterson 2015. We mention again here those whose contributions were invaluable to this work. Thank you (in alphabetical order): Arbortec Tree Expert Company, J. Dickinson, R. Kimball, I.J. Lovette, H. Moravec, P. O’Malley, S.C.R. Smith, L.M. Stenzler, A.K. Townsend, R. Van Den Bussche, and D.M. Woods. We are also very thankful for the unconditional support and cooperation we received from Stillwater residents. A couple of anonymous reviewers of previous drafts of this work made helpful comments, and we thank them, too.

This study was carried out in strict accordance with the recommendations in the Guidelines to the use of Wild Birds in Research of The Ornithological Council, and was approved by the Oklahoma State University IACUC (ACUP No: AS50713). Capture and marking of crows occurred under U.S. Department of Interior, U.S. Geological Society, Federal Bird Banding Permit #22165 (Caffrey). Our work was funded in part by grants from The Payne County Audubon Society, and generous assistance from J. and N. Wilhm, and E. and H. Caffrey.

## APPENDIX

1. Hatching Notes

a. Upon returning to nests after breaks during incubation, females tended to get in and immediately sit, without looking down (below them). Once hatching had begun, when females returned after most breaks, they now looked down into nests, for periods ranging from a few seconds to two minutes, before settling back in.

b. Because hatching is asynchronous, nestling food demand at crow nests starts low and increases as hatching proceeds; nestling-related activities at most nests begin low as well. At several nests of groups with auxiliaries in Stillwater over the years 1997-2000, and three in 2001 and 2002, however, something akin to “excitement” was detectable on presumed hatching days as auxiliaries arrived, sometimes jostling with others, to nudge females and look underneath.

c. Two visits by extra-group individuals on the first days of hatching were observed, both at nests of AM (the female breeder of Group 29 in 2001 and 92 in 2002; below, 3b and 4, and Appendix A 1a in Caffrey and Peterson 2015). In 2001, ZO (a half-sib of at least some of her offspring [Appendix A 1a in Caffrey and Peterson 2015] and who had been snapped at as he approached her during incubation [Caffrey et al. 2016a, Results, Incubation]) was snapped at as he landed on the rim (he left). In 2002, NK, the male breeder of Group 42 and the son of AM and her current (and former) mate (Appendix A 1a in Caffrey and Peterson 2015), landed on the rim of their nest and looked in for several seconds while both were temporarily away.

2. Extra-group individuals making nestling feeding trips

a. BO: Appendix A 3b in Caffrey and Peterson 2015.

b. SV: Appendix A 1b and 3f2 in Caffrey and Peterson 2015.

c. RG: Appendix A 1b and 3b in Caffrey and Peterson 2015.

d. KP: Appendix A 1b and 3e in Caffrey and Peterson 2015.

3. Nestling stage notes

a. In addition to food carried in esophageal pouches, we saw mulberries, pecans, a small white egg (after having been dunked in water), the contents of two separate avian eggs (after being slurped out; the eggs had been carried to the ground and poked open), pieces of a small mammalian carcass stashed in a nearby branch crotch, a nestling American Robin (*Turdus migratorious*), spiders, large insects (legs and wings extended outside crow bills), worms, snakes, and a lizard delivered to nests. When local sources were available – puddles and bird baths – crows would frequently add water to food before delivering to nestlings (and incubating females).

b. In 2001, the nestlings in Nest 29 were about two weeks old when the group’s one-year old female auxiliary (who had the second-highest nestling feeding rate of all auxiliaries in 2001 [and 2002]) was killed by a Great Horned Owl (*Bubo virginianus*; Appendix A 1a in Caffrey and Peterson 2015). A day or two later, the male breeder was hit and critically injured by a car; he remained alive but unable to get to his nest, and impossible to catch, for almost two weeks. The female breeder (AM; Appendix A 1a in Caffrey and Peterson 2015) fed nestlings on her own for almost two weeks before we provided food (we were unable to observe this group for most of this period [Methods, Background] and had not viewed existing tapes immediately, and so were unaware of the situation; AM was thereafter supplied daily with large amounts of dry and canned cat food, scrambled eggs, nuts, fruit, and table scraps). While on her own and before we provided food, AM more than tripled her previous feeding rate and made more trips to the nest per hour during those days of the nestling period than almost every other individual in our population (Fig. 1a), bringing her individual feeding rate to the average nest-level feeding rate of the rest of the population (Fig. 1b). During the single watch (=4.5 hours) after we provided food and before some of her nestlings (=4) fledged, she seemingly chose to maintain the minimum feeding effort level she had already decided upon (Fig. 1a and b) and to invest the additional energy resources in her own self-maintenance.

c. After six days during which the male breeder (Group 22, 2001; Table 1) had made 26% of 51 feeding trips, the female 29%, and the single auxiliary (an adult social son of both breeders) 45% (this individual fed at greater rates than any other auxiliary in the two years), the male breeder disappeared. The female breeder and auxiliary dropped their previous higher-than-average feeding rates (Fig. 5b) and split the subsequent 41 trips 44% and 56%, respectively. A heavy storm was associated with the failure of this nest 19 days post-hatching. The day following the storm, both the female breeder and auxiliary were observed tending to a single, dying nestling; the female for 37 minutes at one point.

d. Nest 333 in 2001 was situated such that, for a variable number of feeding trips per watch, either the two breeders (DZ and SW) or two of the auxiliaries (BL and OM) could not be distinguished. We did not include any watches in our individual-level analyses wherein more than 19% of nestling feeding trips could not be attributed to individuals (n=7), and for included watches (n=8), we dropped the few trips by unidentified individuals from individual-level analyses.

After 25 days of nestling feeding, a three-year old female (BL, a social “Uncertain” for whom we did not have DNA), who had made 13 (21%) of 62 total feeding trips, disappeared. We did six feeding watches after her disappearance, but on five of them the two breeders could not be reliably distinguished on several trips, and so we plot (Fig. 5c) the combined data for breeders and auxiliaries as well as all available pre- and post-disappearance data for individuals. During the single post-disappearance watch wherein all individuals could be identified (=90 min, Day 32), DZ and SW fed at lower rates than their mean rates prior to BL’s disappearance (DZ made zero feeding trips and SW made two), but otherwise their combined feeding rates remained the same. The two remaining auxiliaries maintained their low rates and as such, the contribution to nestling feeding coming from auxiliaries was lower after BL’s disappearance than before.

e. The male breeder in Group 333, 2002 (Table 1, and Appendix A 1b in Caffrey and Peterson 2015; SW) had made 67% of 15 feeding trips before he disappeared after five days of nestling feeding; his mate (DZ) had made one trip, and one of three auxiliaries (their one-year old son, JY) had made 27% (two auxiliaries, their one-year old daughter [TB] and a two-year old son [VK] of DZ and NK [an auxiliary in the process of budding a territory; Appendix A 1b, and Appendix B Predictions 8 and 12-15, in Caffrey and Peterson 2015], made none). VK had been spending time with both Groups 333 and 55 (that of NK) during nest building and incubation (Appendix A 1b in Caffrey and Peterson 2015), after which Nest 55 was abandoned. He continued to be a member of both groups through fledging (333) and a renesting attempt (55) that we could not monitor, respectively. VK was never observed at Nest 333 until 13 days after hatching and eight days after SW was last seen, after which he made 37% of 117 nestling feeding trips (Fig. 5d). DZ and JY split the seven trips we observed (in 210 minutes) before VK began feeding nestlings 71% and 29%, respectively; thereafter DZ increased her feeding rate and JY reduced his (Fig. 5d). TB was never observed to make a nestling feeding trip.

(In 2001, although regularly observed with other group members away from Nest 333 and often in the general nest area, VK was seen to make only one brief visit to the nest and was never observed to feed nestlings.)

f. Male breeders varied widely in feeding contributions: one male breeder was observed to make only 11 of 110 (1%) of the nestling feeding trips in his group (the female breeder made 55% and one auxiliary 29%), one (with an injured leg; UB in Group 19 [2002], Appendix A 2g in Caffrey and Peterson 2015) made 4 of 62 feeding trips (6.5%; the female breeder made 39% and one auxiliary 32% [another auxiliary – with a broken leg – made 19%]), and one made 8 of 64 (12.5%; the female breeder made 41% and one auxiliary 38%), whereas one made 43 of 71 trips (61%; no auxiliaries), one 75 of 117 (64%; no auxiliaries), and one made 64 of 92 (70%; one auxiliary). Female breeders were also highly variable: one made only 10 of 81 trips (12%; the male breeder made 20% and one auxiliary 47%) whereas another made 22 of 33 (67%; her mate was injured [UB in 2001; Appendix A 2g in Caffrey and Peterson 2015] and her two auxiliaries each made only one trip), and another made 34 of 50 (68%; no auxiliaries).

g. In one case of a female breeder intervening in her mate’s feeding of nestlings, she (AM: above, 3b, and below, 4 [and Appendix A 1a in Caffrey and Peterson 2015], observed on videotape) appeared frenzied when he arrived at the nest (in complete contrast to her reactions to the arrival of her one-year old daughter), and almost always flew in to join him if she was not at the nest when he approached. He put large amounts of food into nestling mouths and AM continually removed and redistributed it. Several times she *took* the food from the male as he landed on the rim; once resulting in a brief tug-of-war before he seemingly acquiesced.

4. AM was the female breeder in Groups 11, 29, and 92 (above, 3b, and Appendix A 1a in Caffrey and Peterson 2015). In 2000, AM and her mate (MZ) nested in a tall pine across the street from one of us (TWH). As we assembled to prepare to climb to her nest (to mark nestlings), AM and MZ became agitated (they were vocalizing, flicking wings and tails, and moving around in trees). They swooped at the tree climber and, as described in Caffrey (2001), AM intentionally broke off pine cones and deliberately dropped them on the climber’s head. Four nestlings were marked and returned to the nest. Six days later, three of the four fledged successfully into trees across the street from the nest tree, and bordering the yard of TWH. A week later, the fourth nestling (TG) finally fledged and was found on the ground in TWH’s yard. From experience, we know that weak fledglings on the ground in areas dense with mammalian predators do not survive. In this case, we decided to try to rescue TG, meaning TWH had to catch him and get him up off the ground and into some vegetation; a process involving chasing a mobile, partly volant, squawking animal. AM, extremely upset, repeatedly gave harsh alarm calls and dive-bombed TWH, once contacting her head. After having to repeat the process twice more (catch TG and place him in supportive vegetation) - because he was weak and unable to maintain his position (he kept falling to the ground) - TWH took TG into her house (we were going to attempt to rehabilitate him). For the next four days, until TWH moved out (an independent event), AM stalked TWH inside her house: AM would perch with a view of the room TWH currently occupied, and would move from window to window as TWH changed locations. AM would sometimes issue harsh alarm calls as she peered through at TWH, but at other times was eerily silent. Ten days after removing TG from the field, one of us (CC) released him in the presence of AM, MZ, and two of TG’s broodmates. As CC retreated, AM and MZ immediately joined TG and seemingly encouraged him to join his siblings (by producing quiet vocalizations [of a type also associated with nestlings; CC, pers. obs.] and flying from him to his siblings and back). AM was preening TG when CC left. We never saw TG again.

